# High resolution analysis of population structure using rare variants

**DOI:** 10.1101/2025.07.18.665597

**Authors:** Lei Huang, Thiseas C. Lamnidis, Stephan Schiffels

## Abstract

Various statistical methods have been developed to identify population structure from genetic data, including *F*-statistics, which measure the average correlation in allele frequency differences between two pairs of populations. However, the SNPs analyzed with *F*-statistics are often limited to those found as part of microarrays or, in the case of ancient DNA, to SNP capture panels, which are those within the common allele frequency band. Recent advances in sequencing technology increasingly allow generating whole-genome sequencing data, both ancient and modern, which not only enable querying nearly every base of the genome, but also contain numerous rare variants. Rare variants, with their more population-specific distribution, allow detection of population structure with much finer resolution than common variants - an opportunity that has so far been under-exploited. Here, we develop a new statistical method, *RAS* (Rare Allele Sharing), for summarizing rare allele frequency correlations, similar to *F*-statistics but with flexible ascertainment on allele frequencies. We test *RAS* on both published and simulated data and find that *RAS* has better resolution in distinguishing populations, with appropriate ascertainment. Leveraging this, we further develop the use of *RAS* to compute ancestry proportions with higher accuracy than existing methods, in cases of closely-related source populations. We implemented the new statistical methods as an R package and a command line tool. In summary, our method can provide new perspectives to identify and model population structure, allowing us to understand more subtle relationships among populations in the recent human past.

## Introduction

Human population structure is shaped by past demographic events, which in turn can be inferred using genomic data. For example, populations isolated from each other for an extended period of time will differ in their allele frequencies due to genetic drift (exacerbated in case of small population sizes). On the other hand, migrations and admixture tend to equalize allele frequencies and affect population structure. Therefore, by analyzing and modeling population structure we can infer demographic processes. To identify population structure and make inferences on past demographic events (such as isolation and migration), many statistical methods have been developed and established in the field.

One popular approach is *F*-statistics (Patterson et al. 2012; Peter 2016), subdivided into *F*_2_, *F*_3_ and *F*_4_ depending on the number of populations involved. All *F*-statistics can be formulated as *F*_4_, and therefore can measure the average correlation in allele frequency differences between two pairs of populations (Reich et al. 2009; Lipson 2020), and reflect the overlap between two genetic drift paths in a demographic model of the relationships of all populations involved (Reich et al. 2009; Patterson et al. 2012). *F*-statistics were first proposed in Reich *et al*. (2009) to test for “tree-ness” and compute the admixture proportion of focal populations rejecting tree-ness. Hereafter, *F*-statistics have been widely applied in human archaeogenetics, such as testing genetic similarity of populations (Raghavan et al. 2014), determining ancestry components (Reich et al. 2012; Lazaridis *et al*. 2014) and detecting past admixture events (Patterson *et al*. 2012).

One critical advantage of *F*-statistics, unlike many other methods relying on allele frequency modeling (e.g. momi (Kamm *et al*. 2020), fastsimcoal (Excoffier *et al*. 2013), dadi (Gutenkunst *et al*. 2009)), is their robustness to some forms of SNP ascertainment. Specifically, it was shown theoretically (and in simulations) that statistical tests based on *F*-statistics (e.g. for tests for admixture) are unbiased under certain outgroup-directed ascertainment schemes (Patterson *et al*. 2012). It turns out, empirically, that non-outgroup-directed ascertainments are also close to being unbiased (although see Flegontov *et al*. (2023)). The ability to analyze SNP-ascertained datasets was and still is critical, as Array-Genotyping (e.g. Illumina 650K (Li *et al*. 2008), Affymetrix Human Origins (Patterson *et al*. 2012), 1240K (Mathieson *et al*. 2015)) remain the primary tool for obtaining genome-wide variation for population structure analysis including *F*-statistics (Novembre and Stephens 2008; Li *et al*. 2008; Patterson *et al*. 2012). In the mean time sequencing cost dropped further and larger amounts of whole-genome sequencing datasets were generated in recent years (The 1000 Genomes Project Consortium 2015; Bergström *et al*. 2020). In the past several years we have also witnessed an explosion of ancient DNA data, in large part based on enrichment techniques on ascertained SNP sets (Haak *et al*. 2015; Mathieson *et al*. 2015), but increasingly also based on shotgun sequencing, due to improvements in extraction methods and sequencing technology (Orlando *et al*. 2021).

The ongoing shift to shotgun sequencing, both in modern and ancient DNA analysis, makes it possible to querying nearly every base of the genome, allowing the application of advanced demographic inference methods, as well as addressing potential bias in *F*-statistics of samples genotyped via SNP arrays or insolution target capture (Flegontov et al. 2023). More importantly, whole-genome sequencing data contain many rare variants (The 1000 Genomes Project Consortium 2015; Bergström et al. 2020), which are more likely to be recently derived and can lead to novel conclusions on recent demographic history. For example, signals of recent admixture can be emphasized, as different African populations share more doubletons (variants shared by only two individuals, i.e with allele count two) with each other than with East Asians, which is also reflected in the significant positive value of *D*(Chimp, Yoruba; Han, Mbuti) at low derived allele frequency in Yoruba, contradicting the phylogeny (Mbuti, (Yoruba, Han)) inferred from genome-wide variation (Bergström et al. 2020). Rare variation has been recognized as a potential tool for identifying fine-scale population structure, especially for distinguishing closely related populations. When quantifying the affinity between pairs of individuals in the 1000 Genomes Project with doubleton sharing, rather than genotype covariance, boundaries among populations are more pronounced, and even subgroups in populations GBR (from Britain) and CHS (from Southern China) can be detected. That is because many more doubletons are shared within the same subgroup than between different subgroups of the same population (The 1000 Genomes Project Consortium 2015). Rare allele methods have also been successfully applied to ancestry estimation. When focusing on alleles with lower frequency, ancient British populations from the Iron Age and Anglo-Saxon period are better distinguished by the ratio of alleles they share with Dutch and Spanish, therefore allowing accurate estimation on the Anglo-Saxon British ancestry in present-day British populations (Schiffels et al. 2016). Similarly, among North American indigenous groups, present-day Athabaskans can be distinguished from other groups due to recent admixture from Paleo-Eskimos, and therefore can be modeled as being admixed between northern First Peoples and Paleo-Eskimos (Flegontov et al. 2019).

However, rare allele methods have not yet been coherently formalized in earlier publications. Considering the similarity between *F*-statistics and previous rare allele methods, in this article we incorporate rare allele analyses into the definition of *RAS*-statistics, and demonstrate through simulation and empirical data their ability to outperform ordinary *F*-statistics in detecting recent demographic events, even when the latter are applied to whole-genome data. We derive a *RAS*-based method for ancestry decomposition and show that it gives more accurate estimates than *F*-Statistics based ancestry proportions.

## Method

### *RAS*: Rare allele sharing statistics

Here, we define a statistic summarizing rare allele frequency correlations, *RAS*, an acronym for “Rare Allele Sharing”. *RAS*- statistics are computed on genome-wide biallelic SNPs, similar to *F*-statistics (Patterson et al. 2012) but with ascertainment on rare variants.

In order to define *RAS*-statistics, we first define the following concepts:

#### Reference population

*R* A group of individuals that is used to ascertain variants within specific allele frequency ranges.

#### Outgroup

*O* An individual or group, which is an outgroup to all other individuals/groups involved in the analysis. It is used to define the ancestral allele, and hence polarize alleles into ancestral and derived.

#### Genome length

*L* The number of all positions in the genome considered for analysis. Usually this covers all biallelic SNPs in large panels such as 1000 Genomes (The 1000 Genomes Project Consortium 2015) and/or HGDP (Bergström *et al*. 2020).

#### Sample allele frequencies

**x**_*A*_ A vector of length *L* representing outgroup-directed derived allele frequencies in population *A*. We use expression **x**_*A,i*_ to refer to sample allele frequency *A* at position *i*.

#### Ascertained SNP set

*M*(*O, R, f*_**min**_, *f*_**max**_) The set of all positions at which the derived allele frequency (polarized via outgroup *O*) in reference population *R* is between *f*min and *f*_max_.

#### Ascertained non-missing overlap

*L*_*M*_ (*A, B*) the number of sites in *M* that are non-missing in population *A* and *B*.

We then define a simple statistic as the correlation of two frequencies (for brevity, we write *M* instead of *M*(*O, R, f*_min_, *f*_max_)):

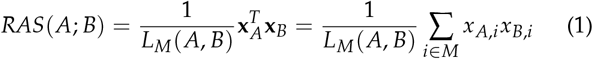

Intuitively, this statistic measures the average rate of allele sharing among any pair of haplotypes from groups *A* and *B* across all ascertained variants.

Indeed, there is a close correspondence of *RAS*(*A*; *B*) and so-called outgroup-*F*_3_-statistics, which are more generally defined as

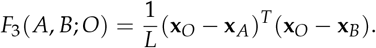

Using derived allele frequencies polarized by our ascertainment outgroup *O*, and using only monomorphic sites in *O*, this definition simplifies to 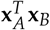 as in equation 1.

From our basic form of *RAS*-Statistics (eq. 1), we derive the following *RAS*-differences, termed *RASD*:

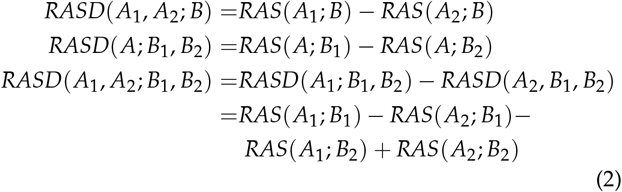

Those derived *RASD*-statistics can be used to test symmetry or treeness, similar to the widely used *F*_4_ or *D* statistics (Patterson *et al*. 2012). *RASD*(*A*_1_, *A*_2_; *B*) corresponds to *F*_4_(*A*_1_, *A*_2_; *B, O*), which tests relative sharing between *A*_1_ and *A*_2_ with respect to *B*. Similarly, *RASD*(*A*; *B*_1_, *B*_2_) corresponds to *F*_4_(*A, O*; *B*_1_, *B*_2_). Ultimately, the difference of differences *RASD*(*A*_1_, *A*_2_; *B*_1_, *B*_2_) corresponds to *F*_4_(*A*_1_, *A*_2_; *B*_1_, *B*_2_).

Note that this correspondence to *F*_4_ statistics becomes an equivalence in the special case that all sites are non-missing in all considered groups, or only sites non-missing in all groups are considered (e.g. using `maxmiss = 1` in the Software `qpfstats` Huang, Lamnidis and Schiffels 3 from *ADMIXTOOLS2* (Maier et al. 2023), and no allele frequency ascertainment is applied (i.e. *f*_min_ = 0 and *f*_max_ = 1). In the more general case relevant here, due to different patterns of missing SNPs in ancient samples, our linear combinations in the definition of *RASD* are not equal to *F*_4_ even without frequency ascertainment.

#### Ancestry decomposition

We can use *RAS* and *RASD* statistics to compute ancestry proportions. Specifically, we model a given *target population T* as a linear sum of *source populations { S*_1_, *S*_2_, …, *S*_*n*_*}*, with the coefficients *{ β*_1_, *β*_2_, …, *β*_*n*_ *}* summing up to 1. Symbolically, we can write

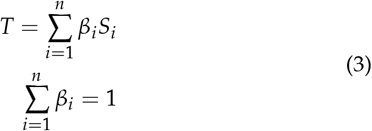

The key idea is to represent the *target* and *sources* by their shared genetic drift (as estimated using *RAS*-statistics) with a selected group of *reference populations*. Following the nomenclature from the popular qpAdm Software (Patterson et al. 2012; Maier et al. 2023), we denote *target* and *sources* as *left populations*, and *references* as *right populations*.

Specifically, we choose a set of *m right populations {R*_1_, *R*_2_, …, *R*_*m*_*}*. If *T* is then admixed as specified in equation 3, and if there was no gene flow going from the *left* into the *right populations* (only from *right* to *left*, see below for a discussion on relaxing this condition), then we can write:

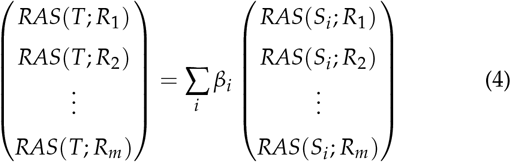

Defining the vectors **v** = *{v*_*j*_*}* = *{RAS*(*T*; *R*_*j*_)*}* and **b** = *{β*_*i*_*}*, and the matrix **W** = *{W*_*ij*_*}* = *{RAS*(*S*_*i*_ ; *R*_*j*_)*}*, we can write this as:

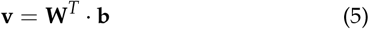

This would be a simple linear regression model, but we still have to satisfy the constraint 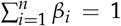. We therefore first write the last element of **b** as

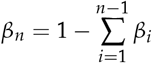

Restricting to a single row *j* of the equation, we then get

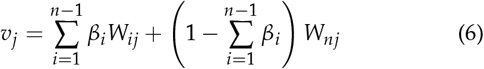

which in turn becomes

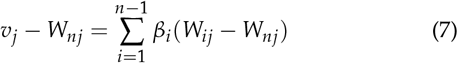

The differences on both sides of the equation are actually *RASD*-statistics. We define *X*_*ij*_ = *W*_*ij*_ *− W*_*nj*_ = *RASD*(*S*_*i*_, *S*_*n*_; *R*_*j*_) for *i* = 1, …, (*n −* 1), *y*_*j*_ = *v*_*j*_ *− W*_*nj*_ = *RASD*(*T, S*_*i*_ ; *R*_*j*_) and **a** = *{β*_1_, …, *β*_*n−*1_ *}* can then write

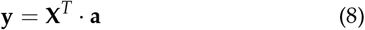

Or

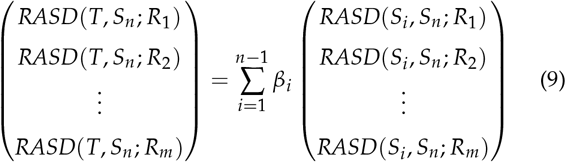

which is a simple linear regression equation. The least-square solution of this equation is (see Hastie *et al*. (2009)):

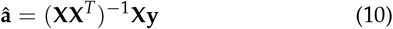

which is a genome-wide point-estimate of admixture proportions. There are two sources of uncertainty/error to consider in this estimation: First, the sampling noise and the finite length of the genome, and second the standard error from the least-square fit itself. We estimate both of these errors using a genome-wide block-Jackknife procedure (Busing *et al*. 1999) to re-estimate the admixture proportions using equation 10 with each of the blocks removed, and derive the standard error of them. This approach is inspired by the Jackknife implemented in qpAdm and the ADMIXTOOLS (Patterson *et al*. 2012) and ADMIXTOOLS2 packages (Maier *et al*. 2023)

The basic decomposition (equation 4) relies on there being no gene flow from left to right populations, only vice versa. This is hardly ever true in real populations. However, in contrast to ordinary F-Statistics, which is the basis for ancestry decomposition in qpAdm, RAS values are only affected by reverse gene flow if it occurred very recently. This can often be ruled out. But even if not, violations of the no-left-to-right gene flow assumption may affect goodness-of-fit statistics more than the actual ancestry estimates which we focus on here.

### Implementation of the method

All scripts used to process and analyze data, exclusively in the R programming language (Ihaka and Gentleman 1996), are provided within a GitHub repository https://github.com/huanglei-artificium/RAS_tools, including documentation.

Briefly, our tool computes *RAS* and *RASD* with flexible ascertainment on allele frequency. Besides .geno files, our tool also accepts allele frequency data consisting of two columns representing numerator (nr of alternative alleles) and denominator (nr of non-missing haplotypes) as input. Then we compute allele frequencies for each population and each site, which are then used to i) select sites that fulfill ascertainment conditions, and ii) to compute the actual *RAS*-statistics on those sites.

To compute optional uncertainties based on a blockwise jack-knife estimate (Busing *et al*. 1999), *RAS* gets computed blockwise (typically by chromosome), which are then combined to obtain genome-wide statistical values.

Multiple statistics, cycling through several populations, and multiple ascertainment conditions are handled efficiently inside our tools, and can be computed in one run.

### Simulations

To illustrate how *RAS* and *RASD* perform in contrast to *F*_3_ and *F*_4_ statistics, we devised a simulation scheme that allows for varying levels of population structure by tuning migration rates. Specifically, we use msprime (Kelleher et al. 2016) to simulate a set of nine populations, located in a 3 *×* 3 grid, each with an effective diploid population size *N*_*e*_ = 20, 000 individuals, with *n* = 50 diploid samples drawn from each population. Each population is connected to its non-diagonal neighboring populations through a symmetric two-way per-generation migration rate. All the migration rates (*m*) in the same simulation were uniform, and varied across different simulations to produce different degrees of population structure among the simulated populations, with the scaled migration rate (4*mN*_*e*_) being 1, 2, 5, 10, 20, 50, 100, 200, 500, 1000, 2000 and 5000 respectively. Each sample has 20 chromosomes, each 100Mbp in length. We number our nine populations from 0 to 8 following a left-to-right, bottom-to-top order, as depicted in Figure 1A. No specific outgroup was simulated, instead we just used a genome consisting exclusively of ancestral alleles as outgroup.

**Figure 1.**
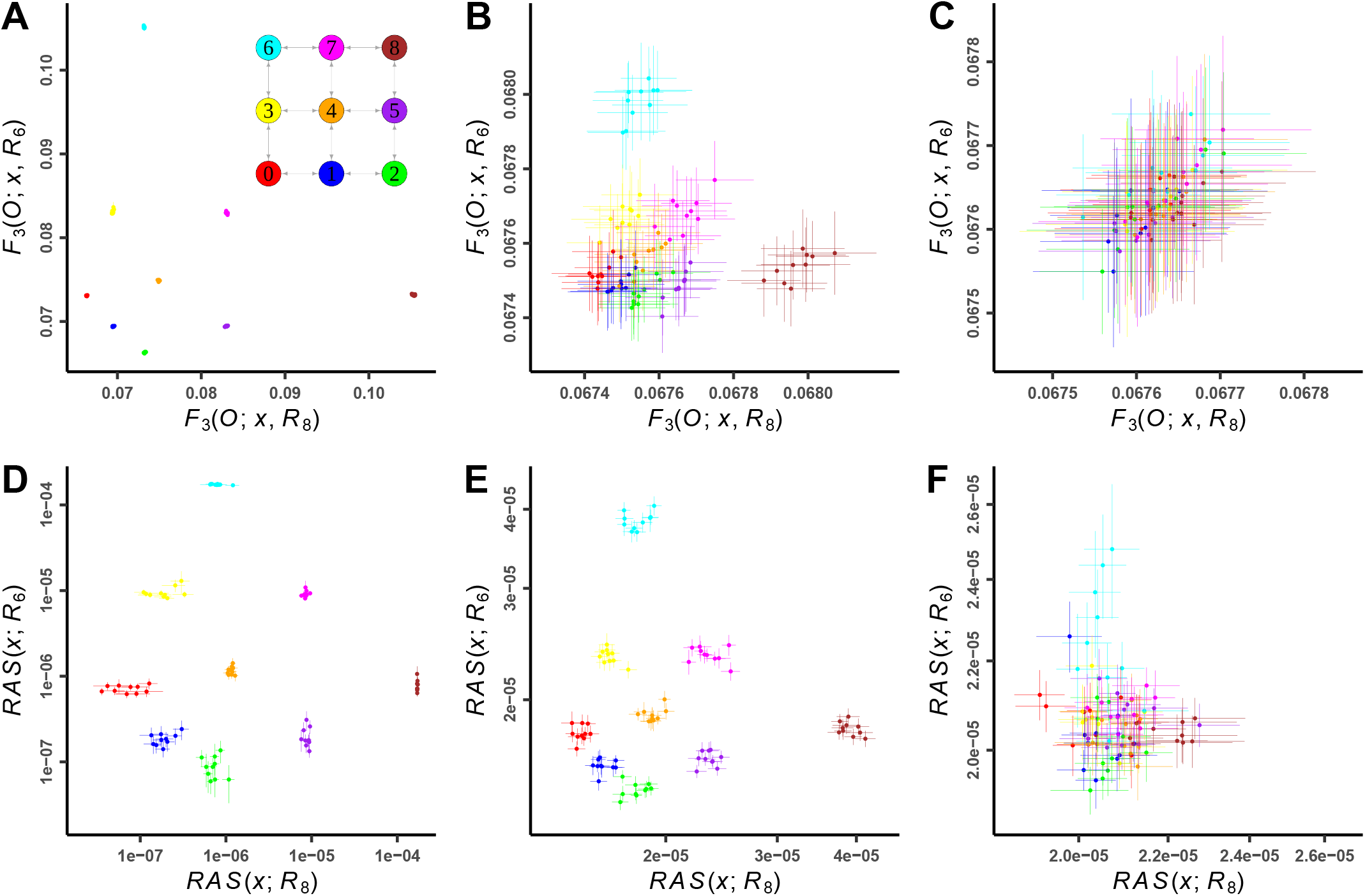
Outgroup-*F*_3_ and *RAS* statistics on test individual *x* and a specific reference population (*R*_6_ or *R*_8_) at different migration rates. The SNP panels used are all sites (A, B and C) and sites with derived allele frequency less than 1% in all reference individuals (D, E and F). The migration rates (4*mN*_*e*_) used in the simulation are 1 (A and D), 100 (B and E) and 2000 (C and F). Test individuals *x* are distinguished by colors representing different populations, shown in legend in (A), which also includes the schematic of simulated population migration. The coordinates of values of *RAS*-statistics are logarithmic.

To assess the population structure in the simulated data, we perform the statistical analysis across three different ascertainment schemes (i) Using all variants in the simulated data, (ii) Using only rare variants with different ascertainment conditions, (iii) Using only a subset of 1.2M variants with derived allele frequency between 0.05 and 0.95, to mimic the 1240K panel (Mathieson et al. 2015).

We chose 10 individuals from each population to act as “test individuals” (denoted by *T*_*i*_), whereas the other 40 individuals are then used as References *R*_*i*_. The ascertainment is therefore based on 9 *×* 40 = 360 reference individuals. For some analysis, in order to average sampling noise, we rotated test and reference individuals 5 times so that each individual served as a test individual once. The code for running the simulation is available at https://github.com/Schiffels-Popgen/RAS_exploration.

### Modern reference data

For our modern reference data, we chose as a starting point the recently released harmonized dataset of 1000 Genomes Project (1kGP) and Human Genome Diversity Project (HGDP) (Koenig et al. 2024), where a new genotype calling was made based on the raw sequencing data from 1kGP and HGDP, with more than 150 million high-quality variants identified, including a large number of rare variants. We chose to focus on the European populations in this dataset, which includes five from 1kGP and eight from HGDP, in which genetic outliers and relatives closer than second-degree were filtered out according to the analysis of Koenig *et al*. (2024) (see Table 1 for the number of individuals for each population). We further supplemented this basic dataset with three European public datasets with genome-wide allele count data: Danish from GenomeDenmark project (Maretty *et al*. 2017), Dutch from Genome of the Netherlands (GoNL) project (The Genome of the Netherlands Consortium 2014) and Swedish from SweGen project (Ameur et al. 2017) (Table 1).

**Table 1.** Number of haploid copies (2*N*) of present-day populations used for analysis

| Population abbr. <sup>a</sup> | Population | Haploids | Dataset |
| --- | --- | --- | --- |
| CEU | Northern and Western European ancestry | 242 | 1kGP |
| FIN | Finnish | 196 | 1kGP |
| GBR | British | 176 | 1kGP |
| IBS | Spanish | 208 | 1kGP |
| TSI | Toscani in Italy | 206 | 1kGP |
| - | Adygei | 34 | HGDP |
| - | Basque | 46 | HGDP |
| - | French | 54 | HGDP |
| - | Italian | 22 | HGDP |
| - | Orcadian | 26 | HGDP |
| - | Russian | 50 | HGDP |
| - | Sardinian | 54 | HGDP |
| - | Tuscan | 16 | HGDP |
| DK | Danish | 40 | GenomeDenmark |
| NL | Dutch | 998 | GoNL |
| SE | Swedish | 2000 | SweGen |
| AFR_all | All Africans | 1978 | 1kGP+HGDP |

<sup>a</sup> Where available, we use abbreviations in the figures.

Even though we focused on Europe, for all the *RAS* statistics regarding real data, we used all African groups in 1kGP and HGDP as outgroups, ascertaining to strictly fixed sites within Africa.

We screened 133 million biallelic SNPs from the harmonized dataset of 1kGP and HGDP for analysis. Variant sites in the Danish, Dutch and Swedish datasets were filtered to this 1kGP+HGDP SNP set, to ensure that our African outgroup is available on all analyzed SNPs. We excluded sites with different alternative alleles when joining the datasets. Unless otherwise stated, we used all European populations in Table 1 for references, which consist of 16 populations and 2184 individuals (4368 sets of haploid chromosomes). Allele frequencies for each site are based on all non-missing individuals.

### Ancient genomes

Our ancient dataset consists of 34 individuals from Great Britain with shotgun-sequencing data, 7 dating to the Late Iron Age (LIA) and 27 to the Early Middle Ages (EMA) (Martiniano *et al*. 2016; Schiffels *et al*. 2016; Gretzinger *et al*. 2022). We started with alignment files (`.bam files`) and called variants overlapping with our reference SNPs, using the Majority-Call method with a minimum coverage of 3 and downsampling in pileupCaller (article submitted). We then selected individuals with more than 1 million SNPs overlapping with our reference data set (see above). We used the ancestry decomposition published in Gretzinger *et al*. (2022), with two major components: CNE (“Continental North European”) and WBI (“Western British-Irish”), and labeled individuals whose dominant ancestry (CNE or WBI) is more than 70% as “England_CNE” (*N* = 17) and “England_WBI” (*N* = 7).

## Results

### Exploring *RAS* with simulated data

We first explored, how our *RAS*-Statistics can distinguish populations from our simulated 3 *×* 3 grid of connected populations (Methods). We chose two corner populations as references, namely population 6 (in the top left, see Figure 1A) and population 8 (in the top right) and computed *RAS*(*x*; *R*_6_) and *RAS*(*x*; *R*_8_) for all test individuals *x* across all populations. For comparison, we also computed *F*_3_(*O*; *x, R*_6_) and *F*_3_(*O*; *x, R*_8_), without any allele frequency ascertainment.

For low migration rate, both *RAS* and *F*_3_ reveal clearly separated clusters of individuals, corresponding to the 9 populations (Figure 1A, with populations closer to the references being placed high on the respective axis, and populations more distant to the references being placed low on the axes (Figure 1A).

Increasing the migration rate shows shows less clearly defined clusters in the case of *F*_3_, with no apparent structure being visible for the highest migration rate tested here (4*mN*_*e*_ = 2000) (1B and C). In contrast, ascertaining SNPs to be rare with respect to the reference populations (which here comprise 9 *×* 40 individuals, excluding the 9 *×* 10 test individuals, see Methods) reveals structure being visible also at higher migration rates. For example, at 4*mN*_*e*_ = 100 (Figure 1E), *RAS* scatter plots reveal still well-separated clusters while *F*_3_ (Figure 1B) already shows considerable overlap between groups. Even at 4*mN*_*e*_ = 2000 (Figure 1F), *RAS* still shows some power to distinguish groups, whereas *F*_3_ appears random (Figure 1C). For our simulated “1240K” dataset with only 1.2 million common variants, structure is substantially less resolved (Supplementary Figure S1). With the highest simulated migration rate, even rare variation appears quite random (Supplementary Figure S1).

Following this qualitative assessment of the ability to separate closely related groups, we devised a more quantitative assessment, by testing for each individual whether it is closest to the mean position of their own population or to some other population, in which case we consider it misclassified. The misclassification ratio is then the proportion of misclassified test individuals relative to all test individuals (using the rotation scheme described in Methods, this amounts to 450 tests). As expected, *RAS* performs better in distinguishing populations for medium and high migration rates, although at the highest migration rate 4*mN*_*e*_ = 5000, the improvement is relatively weak (Supplementary Figure S2). Specifically, at 4*mN*_*e*_ = 50, 100, 200, while *F*_3_ statistics (for both “all sites” and 1240K) start to misclassify, *RAS* can still distinguish the test individuals with no or little error. At low migration rates 4*mN*_*e*_ = 1, 2, note that *RAS* estimates are noisier than the corresponding *F*_3_-estimates, due to fewer shared rare variants: Low levels of migration make it difficult for variants to spread between populations, especially for rare variants between geographically distant populations (e.g., populations 0, 1 and 2 with respect to populations 6 and 8), so that the signal of rare allele sharing between them becomes particularly noisy.

### Application to real data

Turning to real data, we first analyzed present-day European genetic diversity. Specifically, we used the five European populations from the 1000 Genomes Project (1kGP) as references (The 1000 Genomes Project Consortium 2015) and eight European populations from the HGDP project as test individuals (Bergström et al. 2020). We then quantified the affinity between 1kGP European populations and each European HGDP individual with outgroup-*F*_3_ and *RAS* (Figure 2). FIN (Finnish) and IBS (Spanish) from 1KGP are selected as references because they are relatively different geographically and genetically. Both whole genome and 1240K SNP sets reveal differential affinities of the HGDP populations with respect to these references (Figure 2B and C). For example, Russian and Sardinian groups are closest to FIN or IBS references, respectively. Ascertaining on rare allele frequencies in the references with *RAS*, these differences become substantially larger (Figure 2A). In particular, Russian and Basque groups from the HGDP share substantially more rare alleles with either FIN or IBS, indicating recent shared ancestry between Russian and Finnish, and between Basque and Spanish. In addition, there are clearer boundaries for some isolated populations, such as Sardinian and Orcadian. All groups are more clearly separated with *RAS* than with un-ascertained variants.

**Figure 2.**
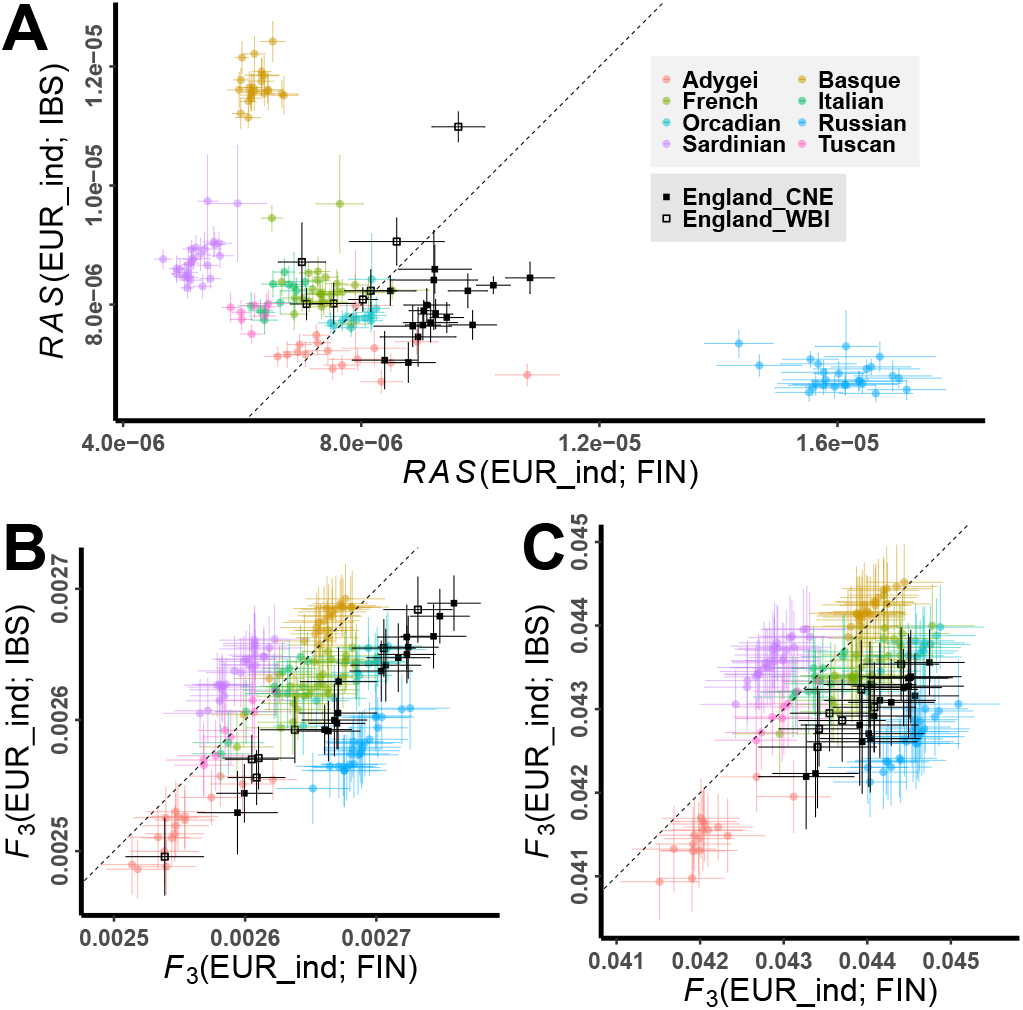
*RAS* and outgroup-*F*_3_ statistics on European individual *x* and FIN/IBS with error bars of ±1 standard deviation (SD). Modern European individuals are distinguished by colors representing different populations in HGDP; ancient British individuals are marked black and distinguished by shapes representing different ancestries. SNP panels are based on the harmonized dataset of 1kGP and HGDP with the following filtering: monomorphic in all 1kGP Africans and derived allele frequency less than 2% in all 1kGP Europeans (A); all sites (B); 1240K sites (C). The dash line indicates the equal relationship to FIN and IBS. FIN: Finnish; IBS: Spanish.

We next turned to ancient DNA to explore the potential of rare variants to analyze ancient population structure. Specifically, we again used the present-day 1kGP reference groups and measured *RAS* and *F*_3_ to a set of ancient genomes from England (Gretzinger *et al*. 2022) for which whole-genome sequencing data is available. Without ascertainment, these two groups do not appear to separate clearly on either all sites or 1240K subset (Figure 2B,C). In particular, WBI individuals are distributed among the entire range of CNE individuals. In contrast, we find that in our RAS analysis (Figure 2A), the WBI individuals fall closer to present-day French compared to samples with CNE ancestry, with all WBI individuals being closer to IBS than to FIN, as indicated by the dashed line in Figure 2A.

### Testing for population structure with *RASD*

A formal test of population structure is a test for symmetry between two groups with respect to a reference group. For classical *F*-Statistics, this is done through *F*_4_-Statistics, which are essentially differences of *F*_3_-Statistics, to statistically quantify differential affinities as deviations from symmetry. Analogously, we defined *RASD*(*A*_1_, *A*_2_; *B*_1_, *B*_2_)-Statistics (Methods), which tests whether *A*_1_ and *A*_2_ are differentially related to *B*_1_ and *B*_2_. Here, we explore the following form:

*RASD*(England_CNE, England_WBI; *R*_1_, *R*_2_), where the first two slots are cycling through ancient individuals from Britain with dominant CNE (*n* = 17) or WBI (*n* = 7) ancestry, respectively, and the last two slots are cycling through various present-day reference populations. We evaluated the results using the Z-Score (i.e. the statistical deviation from zero), using *RASD* at different ascertainment conditions and corresponding *F*_4_ at 1240K and all sites (Figure 3). When applying *RASD*, as long as it is not restricted to extremely rare alleles (up to 0.2%), the ability to distinguish between CNE and WBI is overall better than using *F*_4_, with Z scores even greater than 10 for some combinations. When comparing different reference populations, using NL (Dutch) and SE (Swedish) generally resulted in better discrimination than using FIN (Finnish). This suggests that the immigrants into Britain from continental Europe during Early Middle Age were genetically closer to present-day Dutch or Swedish, consistent with the findings from Gretzinger *et al*. (2022).

**Figure 3.**
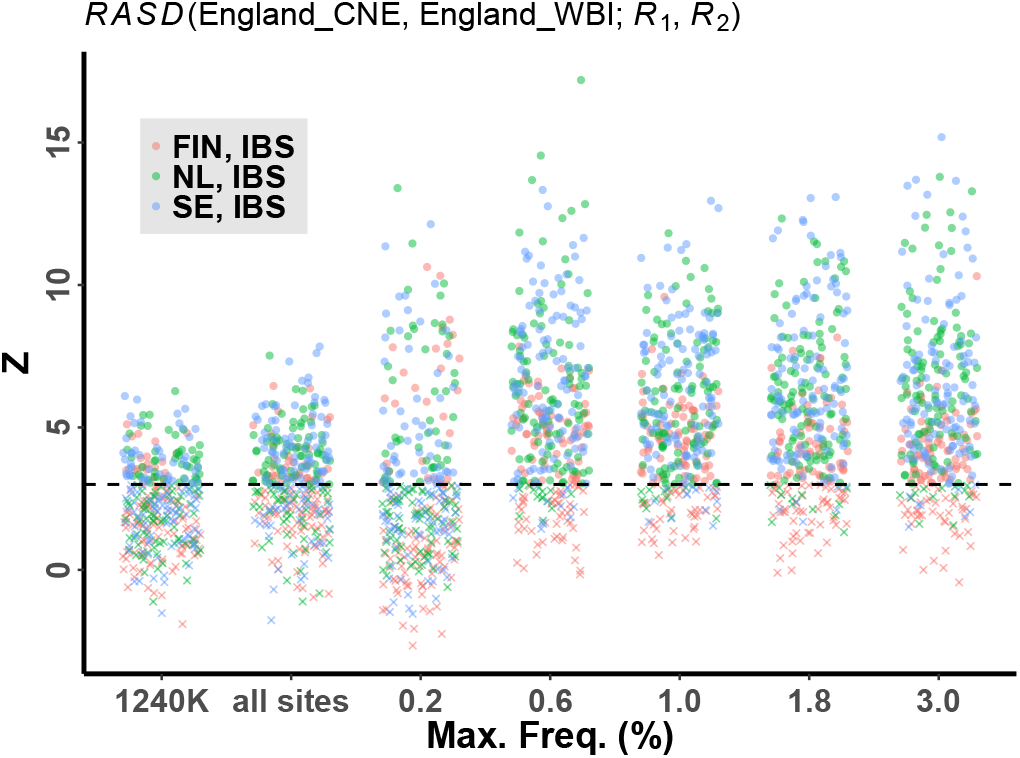
Z score distribution of *RASD*(England_CNE, England_WBI; *R*_1_, *R*_2_) using different SNP panels, shown in each bar with jitter. The ascertained cases are represented in numbers, which are *p*_*max*_ (in percentage) in 1kGP and HGDP European populations plus the Dutch, Danish, and Swedish populations. Different colors represent different reference population pairs (*R*_1_, *R*_2_).

In order to explore the genetic change in Early Middle Age Britain, we performed a systematic analysis by grouping individuals with CNE (*n* = 17) and WBI (*n* = 7) ancestry respectively and comparing them with present-day European populations, under different ascertainment conditions (Figure 4).

**Figure 4.**
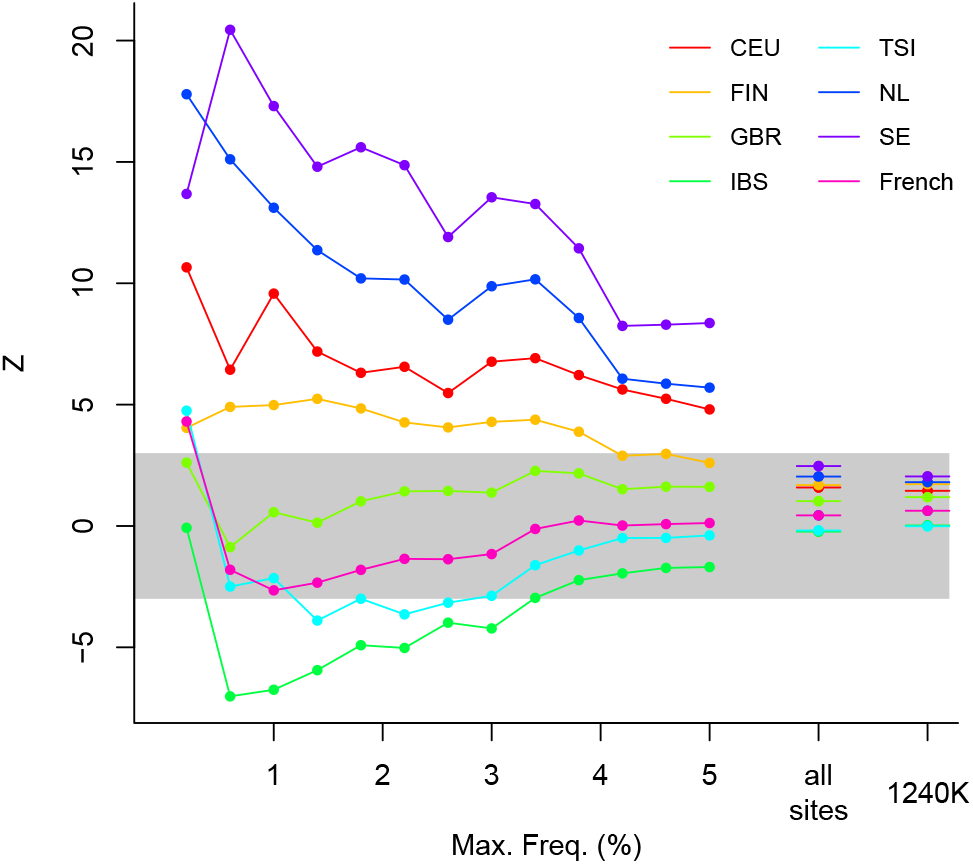
Distinguishing CNE and WBI ancestry using *RASD* and *F*_4_ represented in Z scores. The colored lines are Z scores of *RASD*(England_CNE, England_WBI; *x*), where *x* is a selected present-day European population, under different *p*_*max*_ (in percentage) in present-day European references. Two categories on the right cover ordinary *F*_4_(England_CNE, England_WBI; *x*, AFR_all) on all sites and on 1240K, respectively. The gray area represents |*Z*| *<* 3.

For the 1240K panel and all sites, we observe non-significant positive Z scores for all European reference populations, although some Northern European populations, such as SE (Swedish), NL (Dutch) and FIN (Finnish) have relatively better ability to distinguish CNE and WBI ancestry (Figure 4). In contrast, with *RASD*-statistics, the difference between CNE and WBI, represented by Z score, becomes much more pronounced at low derived allele frequency. Among those present-day European populations, SE (Swedish), NL (Dutch), CEU (Northern and Western European ancestry) and FIN (Finnish) have better ability to distinguish CNE and WBI, suggesting that these present-day populations are most closely related to the actual source that migrated into early medieval Britain, in line with previous conclusion that the immigrants were from Northern Europe (Gretzinger et al. 2022).

Rare alleles can even provide more information about population history at different points in the past, which is reflected by the results of *RASD*-statistics at different derived allele frequency cutoffs. For French, TSI and IBS, their Z scores are much higher at very low frequency 0 - 0.2% than at 0 - 0.6%, which is due to recent low-level gene flow within the European continent making present-day Southern Europeans share increasing number of rare alleles with CNE ancestry, rather than WBI ancestry.

The Z scores of French and TSI at 0 - 0.2% are even slightly higher than those of FIN, reflecting that the magnitude of recent gene flow may be highly dependent on geography, since Finnish and the estimated location of CNE ancestry are on the opposite sides of the Baltic Sea. However, the Z score of FIN reaches its peak at maximum cutoff 1.4% and still remains significant until maximum cutoff 3.8%, suggesting that the gene flow between the CNE ancestry and the ancestor of Finnish occurred in the more distant past, probably mediated through a population closely related to present-day Swedish.

We have compared the ancient British individuals with mostly CNE or WBI ancestry. However, the actual CNE admixture in EMA Britain was like a spectrum, with varying proportions among different individuals (Gretzinger *et al*. 2022). As *F*-statistics are linear under a gradient of admixture components, so are *RAS* and *RASD*. When applied to real data, there is a strong correlation between *RAS* and *RASD*-estimates, and the actual ancestry proportion of a sample. Therefore, we performed *RASD* and *F*-statistics on each EMA British individual by comparing it to a pair of present-day European reference populations that are able to distinguish EMA British individuals. We then computed the correlation between our estimates and the high-resolution CNE ancestry estimated by supervised admixture using thousands of present-day Europeans as reported in Gretzinger *et al*. (2022) (Figure 5; Supplementary Figure S3). At specific *f*_max_, such as 0.6% and 1%, our *RASD* estimates using any of Finnish (FIN), Netherlands (NL) or Swedish (SE) in contrast to Spanish (IBS), show a higher correlation to the reported CNE ancestry estimates than *F*_4_-values on 1240K, indicating a better resolution in distinguishing EMA British individuals with rare alleles, and potentially a more accurate estimation on CNE ancestry from new samples provided the *RASD* statistical values. Those results suggest a potential method for more accurate ancestry decomposition using *RAS*-statistics with appropriate ascertainment on rare alleles.

**Figure 5.**
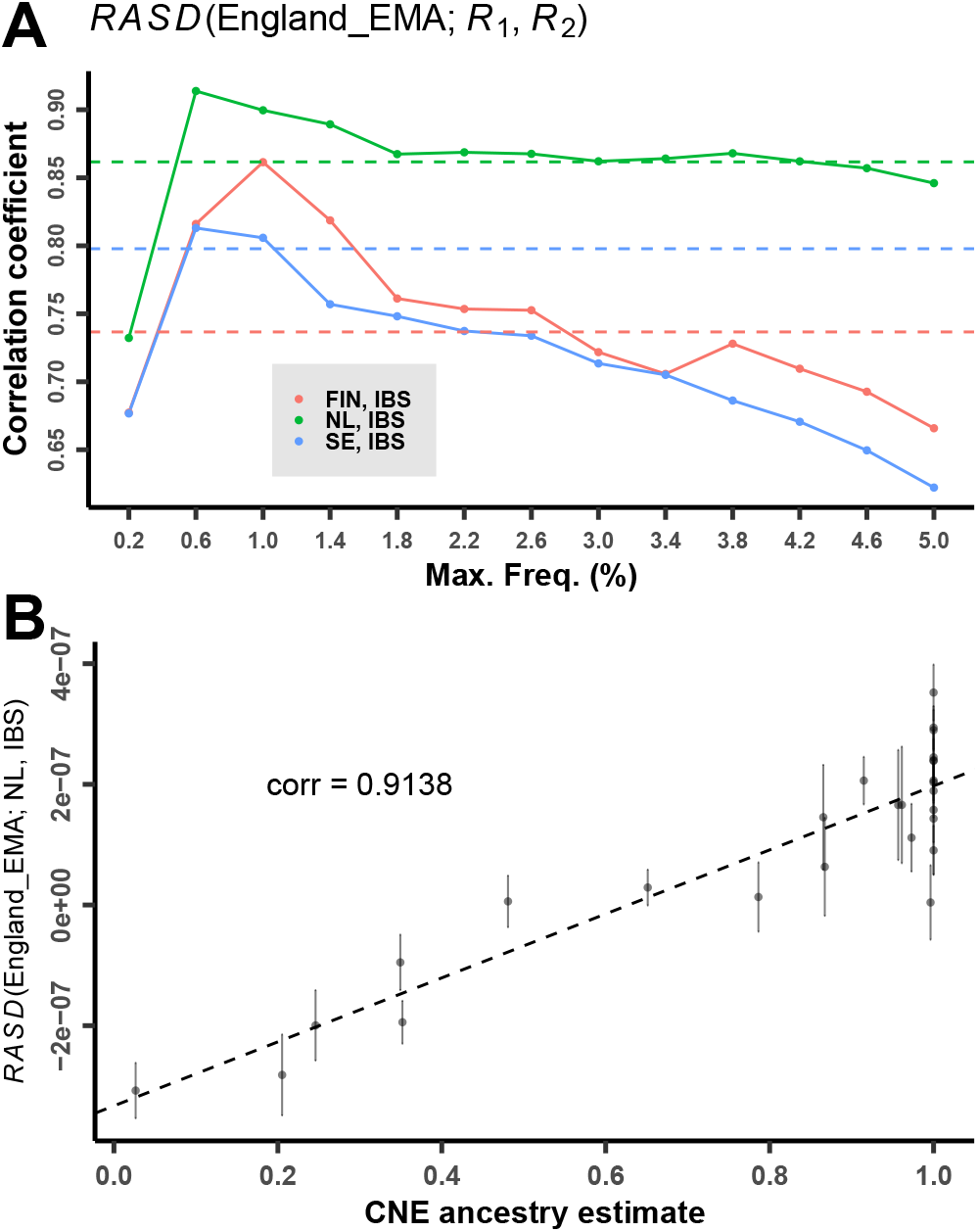
(A) The colored lines are the correlations between *RASD*(England_EMA; *x*_1_, *x*_2_) and estimates of CNE ancestry of each England_EMA individual (from Gretzinger *et al*. (2022)), under different *p*_*max*_ (in percentage) in present-day Europeans. The dashed lines are the correlations of corresponding *F*_4_-statistics on 1240K. (B) *RASD*(England_EMA; NL, IBS) with error bars of ±1 SD, ascertained on sites with 0 – 0.6% derived allele frequency in present-day Europeans.

### Decomposing ancestries using linear combinations of

Motivated by the correlation between *RAS*-Statistics and ancestry components, we devised a new method to decompose ancestry components based on *RAS* (see Methods).

Briefly, every left population (i.e. the target and the sources) has a specific profile of rare allele sharing with each of the right populations, represented by a multi-dimensional vector. We then model the target profile as a linear combination of source profiles, with the coefficient reflecting the admixture proportion.

We tested this new method on our simulated data, focusing on two-component models. According to the simulation scheme (Method), some populations can be represented as admixtures Results with *Z <* 3 are indicated as crosses and with *Z >* 3 with circles.

of other populations. Specifically, we aim to model population 4 (in the middle of the grid) as an admixture of population 0 (bottom-left) and population 8 (top-right). We expect for all migration rates a 50%/50% decomposition, due to symmetry.

We compared our ancestry estimates based on *RAS* with the expected values and defined two types of error: (A) the absolute difference between our estimate and the theoretical value of 0.5, denoted as “true error”; (B) the standard error of our estimate based on a chromosome-wise jackknife (Busing *et al*. 1999), denoted as “self error”.

The results reveal that at high migration rates, both types of errors are substantially lower for our *RAS*-based ancestry estimate compared to *F*_3_-based estimates on 1240K or even on all sites (Figure 6). Note that towards lower migration rates, we observe a turning point (around 4Nm=100, Supplementary Figure S4), where all sites and even 1240K are performing subtly better than rare variants, although at a very low level of error, which we attribute to a lower number of shared rare variants for low migration rates. Therefore in real cases, it’s appropriate to use *F*-statistics when populations are highly differentiated, while *RAS* fills the gap where *F*-statistics lose resolution.

**Figure 6.**
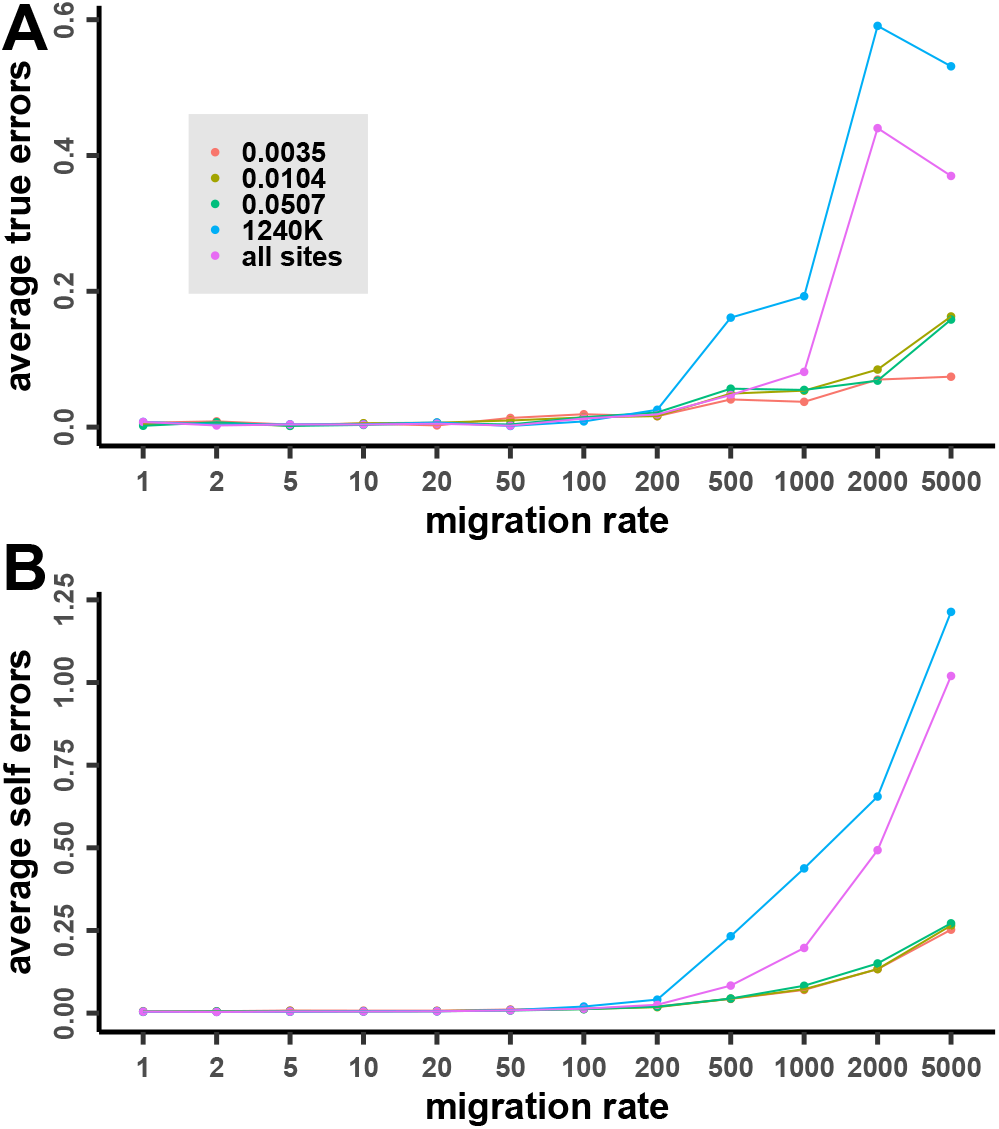
Error estimate for modeling population 4 as an admixture of population 0 and 8. We rotated test individuals for 5 times and calculated the average of errors for 5 parallel tests.

## Discussion

We have defined *RAS*, a statistical method based on rare allele sharing between populations, and demonstrated that rare variation provides powerful means of identifying fine-scale population structure and revealing unique population histories that common alleles may not capture. For both simulated and empirical data, we observe a signal reinforcement of recent demographic events, reflected in the much stronger allele sharing for rare alleles (Figure 1D,E,F; 2A; 3; 4), compared to common alleles and even the whole genome.

Investigating population structure is one of the primary goals of population genetic analyses involving ancient and modern genomes. Such studies have greatly advanced our understanding of historical human populations, and their migration and admixture (Nielsen et al. 2017; Skoglund and Mathieson 2018; Liu et al. 2021; Stoneking et al. 2023). Especially in recent years, studies involving larger sample sizes have revealed more detailed historical demographic events (Lazaridis et al. 2022; Allentoft et al. 2024; Antonio et al. 2024; McColl et al. 2025). In these studies, 1240K remains the primary ascertainment scheme for exploring genome-wide population structure. While increasing sample sizes help with existing methods, we here show that different ascertainment approaches and in particular a focus on rare variants can more dramatically improve resolution, such as the clearly demarcated Russian and Basque populations (Figure 2A). Even for ancient DNA, by embracing an ascertainment scheme strictly in present-day data, the ability to distinguish between populations is also significantly increased, as exemplified here by differentiating between CNE and WBI ancestries (Figure 3;4;5). Surprisingly, the resolution for rare variants is even higher than using all variants (Figure 2;3), which further emphasizes the importance of ascertaining rare alleles for population structure analysis.

Larger sample sizes have a more direct impact on the resolution of our method, compared to most traditional tools based on common alleles such as *F*-statistics and PCA. The rare alleles ascertained from reference populations generally also have low frequencies in the test populations. Therefore, the relative error of their allele frequency estimates is larger than that of common alleles. This highlights the role of increasing sample sizes in improving the accuracy of rare allele frequency estimates, and therefore the resolution of *RAS* statistics. Here, we have compared CNE and WBI ancestries and observed higher *Z* scores when grouping individuals (Figure 4) rather than analyzing them individually (Figure 3). For simulated data, we also group individuals to reduce noise (Figure 6).

In order to get a more complete picture of rare genetic variation, we have used shotgun data for ancient samples. We did not use capture data, due to the far lower amount and uneven distribution of rare alleles, although the raw sequence data still cover some neighboring rare allele sites. Fortunately, recent advances in sequencing technology have made shotgun sequencing more efficient and cost-effective, and shotgun data are becoming increasingly available for ancient DNA studies (Maisano Delser *et al*. 2021; Mallick *et al*. 2024), which coincides with the requirement of *RAS* for a larger sample size. The increasing availability of shotgun data is transforming the field of archaeogenetics as a whole by offering more detailed insights into human population history.

Western Eurasia serves as a prime example: Through the spread of early European farmers from the Near East (Lazaridis *et al*. 2014), and the movement of Indo-European speaking groups from the Eurasian steppe (Haak *et al*. 2015), populations that might have been more genetically distinct have gradually become more similar, leading to an overall more homogeneous genetic structure across Europe, which may have remained stable since the Iron Age (Antonio et al. 2024). Nonetheless, regional differences still persist, which are shaped by local history: during the Roman period, while Northern provinces maintained higher levels of local continuity, Southern sites absorbed the influences from Northern Africa, the Near East, and Eastern European Slavic groups, displaying more genetic variability (Antonio *et al*. 2019; Olalde et al. 2023); Celtic and Germanic tribes occupying different regions of Europe had different genetic profiles due to their different migration routes and interactions with different neighboring groups (McColl et al. 2025). Method development has so far relied on haplotype based analysis, such as IBD-based inference (McColl et al. 2025) or ancestral recombination graph inference (Speidel et al. 2025). In the future, using *RAS*, we will be able to additionally study rare variants that may be overlooked by other methods and expect more insights on subtle demographic events from these comprehensive datasets.

In our final demonstration, we have implemented a new way of estimating ancestry proportions, showing a better performance of *RAS*-based compared to ordinary *F*-statistics-based estimates (Figure 6). In the framework of *F*-statistics, there are extensions based on *F*_4_ matrices: qpWave for testing symmetry or external sources, and qpAdm for testing hypothetical admixture modeling (Patterson et al. 2012). In future work, *RAS*-statistics may be used similarly to develop formal tests for symmetry and admixture.

## Acknowledgments

This project has received funding from the European Research Council (ERC) under the European Union’s Horizon 2020 research and innovation programme (grant agreement number 851511).

## Data Availability

All data used in this manuscript can be accessed from public resources. All modern genomes or allele frequency data (.vcf files) can be obtained from the following websites: the harmonized 1kGP+HGDP dataset at https://gnomad.broadinstitute.org/downloads#v3-hgdp-1kg (Koenig *et al*. 2024), allele frequency data from the GenomeDenmark project at https://ega-archive.org/studies/EGAS00001002108 (Maretty *et al*. 2017), allele frequency data from the Genome of the Netherlands (GoNL) project at https://www.nlgenome.nl/menu/main/app-go-nl/download-data (The Genome of the Netherlands Consortium 2014), allele frequency data from the SweGen project at https://ega-archive.org/studies/EGAS50000000906 (Ameur *et al*. 2017). All ancient genomic data from Great Britain (.bam files or raw read data from the European Nucleotide Archive) from Martiniano *et al*. (2016); Schiffels *et al*. (2016); Gretzinger *et al*. (2022) as described in these publications.

## Code Availability

Software to estimate *RAS, RASD* and ancestry decomposition, as well as to prepare the datasets, can be found at https://github.com/huanglei-artificium/RAS_tools. Documentation on how to use the software is available in the accompanying README file. The scripts used for data simulation are available at https://github.com/Schiffels-Popgen/RAS_exploration.

## Supplementary Figure Captions

Figure S1: Outgroup-*F*_3_ statistics on test individual *x* and a specific reference population (*R*_6_ or *R*_8_) at different migration rates on 1240K SNP panel. The migration rates (4*mN*_*e*_) used in the simulation are 1 (A), 100 (B) and 2000 (C). Test individuals *x* are distinguished by colors representing different populations, shown in legend in (A), which also includes the schematic of simulated population migration.

Figure S2: The misclassification rate for different migration rates.

Figure S3: *RASD*(England_EMA; FIN, IBS) (A) and *RASD*(England_EMA; SE, IBS) (B) with error bars of ±1 SD, ascertained on sites with 0 – 0.6% derived allele frequency in present-day Europeans.

Figure S4: Error estimate for modeling population 4 as an admixture of population 0 and 8. We rotated test individuals for 5 times and did 5 parallel tests. The coordinates of errors are logarithmic.

